# PRSS56 is required for the developmental positioning of ocular angle structures

**DOI:** 10.1101/360321

**Authors:** Cassandre Labelle-Dumais, Nicholas G Tolman, Seyyedhassan Paylakhi, Simon WM John, K Saidas Nair

## Abstract

Angle-closure glaucoma (ACG) is a severe form of glaucoma affecting up to 16 million people worldwide. In ACG, physical blockage of the ocular drainage tissue by the peripheral iris impedes the drainage of aqueous humor resulting in elevated intraocular pressure (IOP) and subsequent optic nerve damage. Despite the high prevalence of ACG, the precise mechanism(s) underlying pathogenesis are only partially understood. We have previously demonstrated that a mutation in the gene encoding the serine protease PRSS56 causes an ACG phenotype in mice. Notably, *Prss56* mutant mice exhibit a reduced ocular axial length and a lens occupying a larger ocular volume compared to WT mice, recapitulating characteristic features of human AGC. Our findings utilizing mouse genetic models demonstrate that loss of PRSS56 function results in altered configuration of ocular angle structures characterized by a posterior shift in the positioning of the ocular drainage tissue relative to the ciliary body and iris during development, leading to a physical blockage of drainage structure (angle closure) and high IOP. Utilizing a previously employed genetic strategy of rescuing mutant *Prss56* mediated reduction in ocular size by inactivation of EGR1 (*Egr1;Prss56* double mutants) we determined the influence of ocular size on developmental positioning of the ocular angle tissues. Our findings suggest that abnormal positioning of the drainage structure as a result of loss of PRSS56 function is uncoupled from its effect on ocular axial length reduction. Furthermore, we demonstrate that the IOP elevation observed in *Prss56* mutant mice is genetic context-dependent and identify a dominant modifier locus on Chromosome 2 of the C3H/HeJ genome conferring susceptibility to high IOP. Overall, our findings reveal a novel role for PRSS56 in the proper configuration of the iridocorneal angle and provide new insight into the developmental pathways implicated in glaucoma pathogenesis.

## Introduction

Glaucoma is a leading cause of blindness and refers to a heterogeneous group of diseases characterized by progressive loss of retinal ganglion cells (RGCs) and corresponding visual field defects. One of the major risk factors for developing glaucoma is elevated intraocular pressure (IOP)^1^. Regulation of IOP mainly relies on the balance between aqueous humor (AqH) production in the ciliary body and its exit through ocular drainage structures located at the iridocorneal angle that forms at the junction between the iris and the cornea^2,3^. The key ocular drainage structures consist of the trabecular meshwork (TM) and the Schlemm’s canal. AqH enters the TM and subsequently drains into the Schlemm’s canal before entering the collector channels, which culminate in the episcleral veins^4,5^.

Consistent with a critical role for the ocular drainage structures in AqH outflow regulation, structural or functional alterations of these structures impair AqH outflow, leading to elevated IOP^6,7^. Depending on the configuration of the iridocorneal angle, glaucoma is broadly divided into primary open angle glaucoma (POAG) and primary angle closure glaucoma (PACG)^7,8^. In PACG, altered positioning of the peripheral iris causes blockade and closure of the iridocorneal angle preventing access of AqH to the drainage tissues. Pupillary block and plateau iris configuration (PIC) are the two major conditions implicated in the etiology of PACG ^7,9,10^. Anatomical features such as reduced ocular size, relatively larger lens, and a shallow anterior chamber predispose individuals to pupillary block, thereby restricting the flow of AqH from the posterior to anterior chamber^11^. The accumulation of the AqH in the posterior chamber causes the iris to bulge anteriorly, leading to its apposition against the ocular drainage tissue. Plateau iris configuration is characterized by the forward displacement of the peripheral iris by an anteriorly localized ciliary body/pars plana that can lead to angle closure^12,13^.

ACG is a complex disease resulting from an intricate interplay between genetic and environmental factors. Although specific anatomical features are well established to predispose to ACG, they are generally not sufficient to induce angle closure/high IOP and other contributing factors remain largely unknown^11^. We have previously demonstrated that mutations in the gene coding for the secreted serine protease PRSS56 result in severe reduction in ocular axial length (nanophthalmos) in humans and many of these individuals go on to develop ACG^14-16^. Consistent with observations in patients, we have recently shown that *Prss56* mutant mice exhibit reduced ocular axial length and develop an ocular phenotype resembling ACG^14^. Of note, their lens diameter is indistinguishable from that of the control eyes, causing the lens to occupy a relatively larger volume in *Prss56* mutant eyes^14,15^. We have previously proposed that these anatomical features could contribute to the ocular angle closure phenotype and IOP elevation observed in *Prss56* mutant mice^14^. It is unclear whether the ACG/high IOP phenotype caused by *Prss56* mutations is a direct consequence of ocular size reduction or arise from additional pathomechanism(s) that are yet to be characterized.

Here, we use a combination of genetic, histological and physiological approaches to determine factors besides reduced ocular size that contributes to the ACG-relevant phenotypes in *Prss56* mutant mice. We have detected altered configuration of the ocular angle structures in *Prss56* mutant mice that is characterized by posterior displacement of the ocular drainage tissues relative to the peripheral iris, leading to angle closure. Our results suggest that the mutant PRSS56 induced misalignment of the ocular drainage structure is uncoupled from its effect on ocular axial length reduction. Finally, we identify a genetic modifier locus that confers susceptibility to high IOP in *Prss56* mutants. Collectively, our findings unveil a novel role for PRSS56 in the developmental configuration of the iridocorneal angle and provide new insight into the pathogenesis of ACG.

## Results

### Loss of PRSS56 function contributes to angle closure and high IOP

We have recently demonstrated that mice homozygous for a null allele of *Prss56* (*Prss56^-/-^)^15^* exhibit ocular axial length reduction, an anatomical feature predisposing to ACG (Fig. 1A). Here, we performed a detailed histological characterization of the ocular angle structures in *Prss56^-/-^* mice in a C57BL/6J (B6) background. In both *Prss56^+/+^* and *Prss56^+/-^* eyes, the drainage structures (TM and Schlemm’s canal) are located anterior to the very deep angle recess (crowded by the iris root and ciliary body) to allow AqH to access the drainage tissues without obstruction by the iris (Fig. 1B). In contrast, the TM and Schlemm’s canal are found in close proximity to the peripheral retina in *Prss56^-/-^* eyes and exhibit a posterior shift in their position relative to the ciliary body and iris (Fig. 1B). Consistent with this observation, the distance between the edge of the peripheral retina and the posterior end of the Schlemm’s canal was significantly smaller in *Prss56^-/-^* eyes compared to *Prss56^+/-^* eyes. Thus, the peripheral iris in *Prss56^-/-^* mice physically obstruct the ocular drainage tissue and resulting in angle closure (Fig. 1 B, C), and develop significantly higher IOP compared to their *Prss56^+/-^* littermates at both ages examined (3 and 5 months, Fig.1D). Together, these findings demonstrate that, in addition to its role in ocular size determination, PRSS56 is required for proper configuration of the iridocorneal angle and that loss of PRSS56 function in the mouse leads to a phenotype reminiscent of human angle closure.

**Figure 1.**
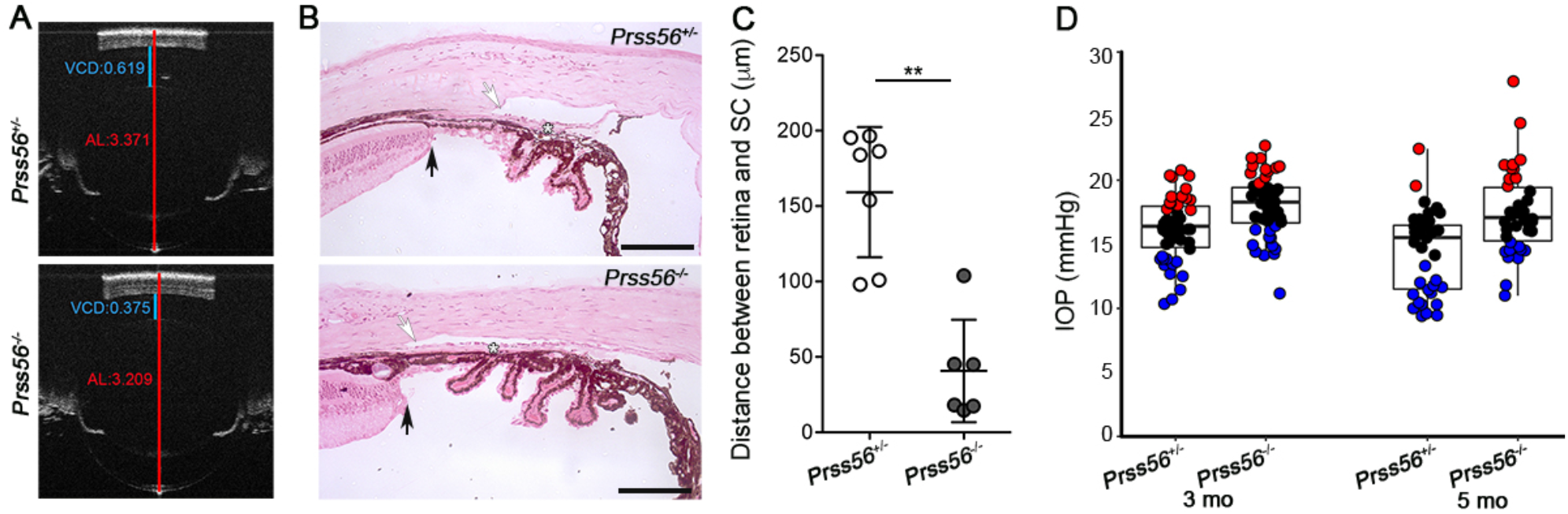
*Prss56* knockout mice show altered positioning of ocular angle structures and develop elevated IOP. (**A**) Representative OCT images showing reduced ocular size in *Prss56^-/-^* compared to *Prss56^+/-^* mice (shown are eyes from 2 months old mice). The red and blue lines indicate ocular axial length (AL) and vitreous chamber depth (VCD), respectively. (**B**) The ocular angle structure of the *Prss56^-/-^* mice show altered organization as compared to the control *Prss56^+/-^* mice. The trabecular meshwork (asterisk) and adjoining Schlemm’s Canal (white arrows) show a posterior shift in their position in relation to the iris and ciliary body, resulting in angle closure. Scale bar=100μm. (**C**) The distance between the peripheral edge of the retina and the posterior end of the Schlemm’s canal are presented as scatter plot. The relative distance separating the peripheral retina (black arrows in B) from the Schlemm’s Canal (white arrows in B) is significantly reduced in *Prss56^-/-^* eyes compared to control *Prss56^+/-^* eyes. Values are presented as mean +/-SD, ** p<0.01, Mann-Whitney test. (**D**) IOP values at ages 3 and 5 months (MO) represented as *k*-means clustered (*k*=3) scatter plots showing significantly higher IOP in *Prss56^-/-^* mice compared to control *Prss56^+/-^* mice. Values are presented as median box plots, N≥ 30 mice per genotype. Similar proportions of males and females were included in each of the experimental group.

### Independent roles for PRSS56 in ocular size determination and iridocorneal angle configuration

The ocular axial length reduction and decreased vitreous chamber depth (VCD) observed in *Prss56^-/-^* mice^15^ may directly contribute to the altered positioning of the ocular drainage structures and elevated IOP. To test this possibility, we took advantage of the *Egr1; Prss56* double mutant (*Egr1^-/-^; Prss56^-/-^*) mouse model we have described recently^15^. In these mice, inactivation of *Egr1* was found to partially rescue the ocular axial length reduction and hyperopia caused by loss of PRSS56 function^15^. Because the ocular axial length of *Egr1; Prss56* double mutants is not detectably different to that of control mice (Fig. 2A)^15^, it allows us to test the contribution of ocular axial length reduction to altered positioning of the drainage structures in *Prss56* mutant mice. Histological analysis revealed a posterior shift in the positioning of drainage structures and angle-closure in *Egr1^-/-^; Prss56^-/-^* mice reminiscent of the altered iridocorneal configuration phenotype observed in *Prss56* single mutants (*Egr1^+/-^; Prss56^-/-^*, Fig. 2 A-B, and Fig 1). In agreement with this observation, the distance between the edge of the peripheral retina and the Schlemm’s canal was significantly smaller in *Egr1^-/-^; Prss56^-/-^* eyes than that of control eyes and comparable to that of *Egr1^+/-^; Prss56^-/-^* eyes. In addition, and consistent with the angle closure phenotype, significant proportion of *Egr1; Prss56* double mutant mice develop high IOP similar to *Prss56* single mutants (Fig 2C). Together, these results demonstrate that *Egr1* inactivation prevents axial length reduction but not angle closure and high IOP in *Prss56* mutant mice. Furthermore, they suggest that the angle closure phenotypes observed in *Prss56* mutant mice is not dependent on ocular axial length reduction, and likely result from altered developmental positioning of ocular drainage structures.

**Figure 2.**
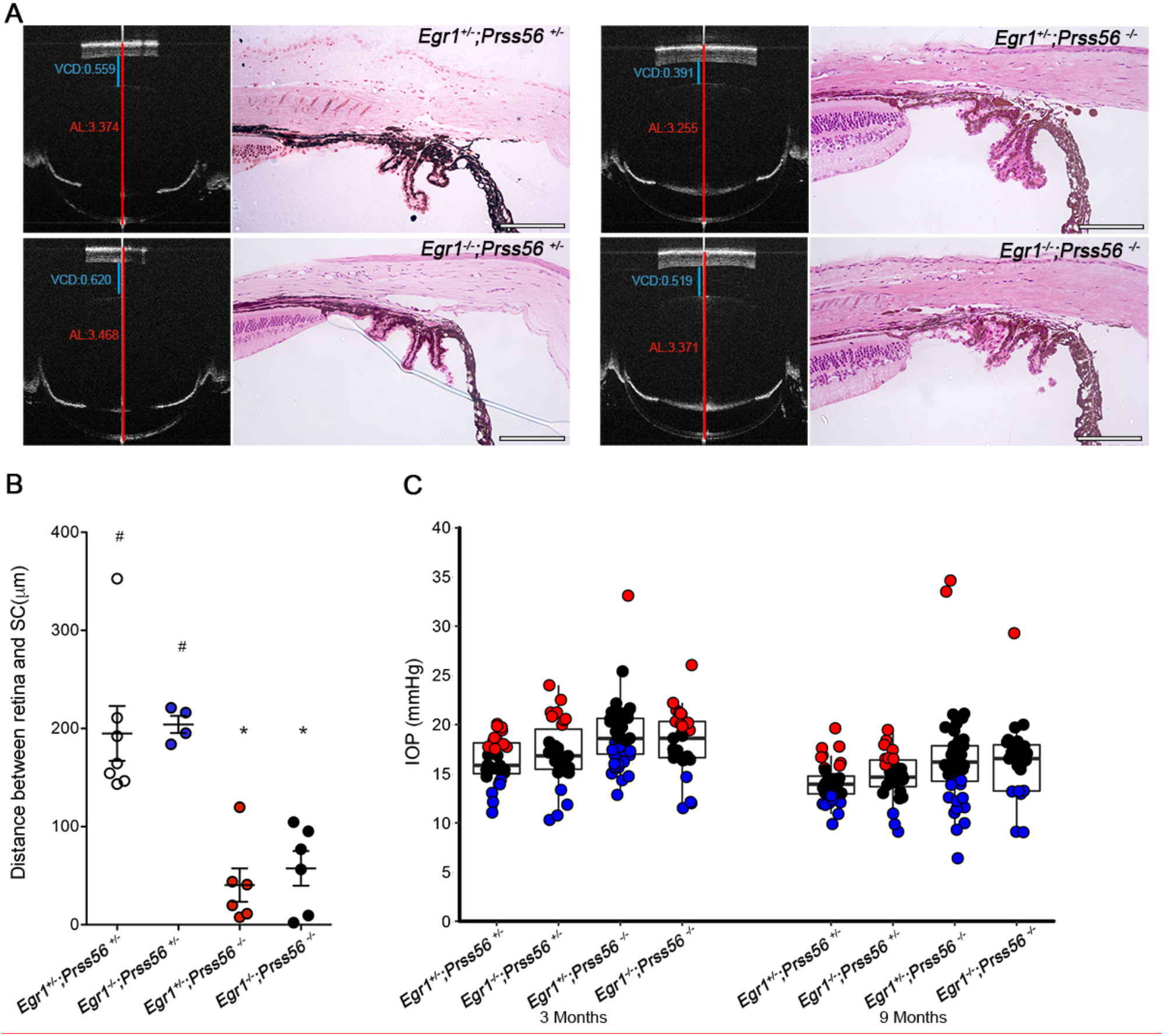
Altered positioning of ocular angle structures is independent of axial length reduction in *Prss56* mutant mice. (**A, B**) Representative OCT images and H&E stained ocular sections demonstrating that *Egr1* inactivation rescues ocular size reduction but not the positioning of the ocular angle structures in *Prss56^-/-^* mice. In OCT images, the red and blue line indicate ocular axial length (AL) and vitreous chamber depth (VCD), respectively. *Egr1* single mutants (*Egr1^-/-^;Prss56^+/-^*) have significantly elongated axial length and *Prss56* single mutants (*Egr1^+/-^;Prss56^-/-^*) exhibit significantly reduced ocular axial length as compared to the control *Egr1^+/-^;Prss56^+/-^ eyes*. In contrast, the ocular axial length of double mutant mice (*Egr1^-/-^;Prss56^-/-^)* is not significantly different from control eyes (*Egr1^+/-^;Prss56^+/-^*). H&E stained ocular sections show a posterior shift of the ocular drainage structure relative to the iris and ciliary body resulting in angle closure in both the *Prss56* single mutants and double mutants (compare *Egr1^+/-^;Prss56^-/-^* and *Egr1^-/-^;Prss56^-/-^* eyes to the controls or *Egr1* single mutants both exhibiting an open angle, shown are 2 months eyes). (**B**) The distance between the peripheral edge of the retina and posterior end of the Schlemm’s canal are shown as scatter plot. The Schlemm’s canal is located closer to the peripheral edge of the retina in both the *Egr1^-/-^;Prss56^-/-^* eyes and *Prss56* single mutants as compared to the control eyes (*Egr1^+/-^;Prss56^+/-^*). Values are presented as mean +/-SEM. For comparison to control eyes (*Egr1^+/-^;Prss56^+/-^*), *p<0.05; for comparison to double mutant eyes (*Egr1^-/-^;Prss56^-/-^*), ^#^p>0.05, Kruskal-Wallis test with Dunn’s multiple comparisons test. (**C**) IOP values from *Prss56* single mutants, *Egr1* single mutants, double mutants and control mice measured at 3 and 9 months are presented as *k*-means clustered (*k*=3) scatter plots. Values are presented as median box plots. The mean IOP value of *Prss56* single mutants (*Egr1^+/-^;Prss56^-/-^)* is significantly higher compared to that of the control eyes at both the ages examined. Interestingly, despite having ocular axial length comparable to that of control eyes the *Egr1^-/-^;Prss56^-/-^* eyes exhibit significantly higher IOP at all ages examined.

### Temporal requirement for PRSS56 for proper configuration of the iridocorneal angle during ocular development

To determine the temporal requirement of PRSS56 for proper positioning of ocular angle structures, we used a conditional gene targeting approach. To this end, we bred conditional *Prss56* mutant mice to the inducible ubiquitous *Ubc-Cre* mouse line to ablate *Prss56* at distinct stages of ocular development. Mice carrying the conditional floxed allele of *Prss56* (*Prss56^F^*) and the inducible *Ubc-Cre* transgene (*Prss56^F/F^*; *Ubc-Cre ER^T2^* or control: *Prss56^F/+^; Ubc-Cre ER^T2^*) were injected with tamoxifen at postnatal day (P) 13 or P18. These time points were selected because the ocular drainage structures are actively developing at P13 and have reached the terminal stage of development by P18 ^18^. Tamoxifen injected animals were aged to P60 for histological analysis of the ocular drainage structures. Following tamoxifen injection at P13 or P18, the control *Prss56^F/+^; Ubc-Cre ER^T2^* eyes were indistinguishable from those of uninjected controls and show an open-angle configuration (Fig. 3B-C). In contrast, altered iridocorneal configuration was observed in *Prss56^F/F^*; *Ubc-Cre ER^T2^* mice injected with tamoxifen at P13 which was characterized by a posterior shift in the positioning of ocular drainage structures similar to that observed in *Prss56*^-/-^ mice (Fig. 1B). When *Prss56^F/F^*; *Ubc-Cre ER^T2^* mice were injected with tamoxifen at P18, the ocular angle structures developed normally and were comparable to those of *Prss56^F/+^; Ubc-Cre ER^T2^* control eyes (Fig. 3C). These findings indicate that PRSS56 is required prior to P18, for proper positioning of angle structures.

**Figure 3.**
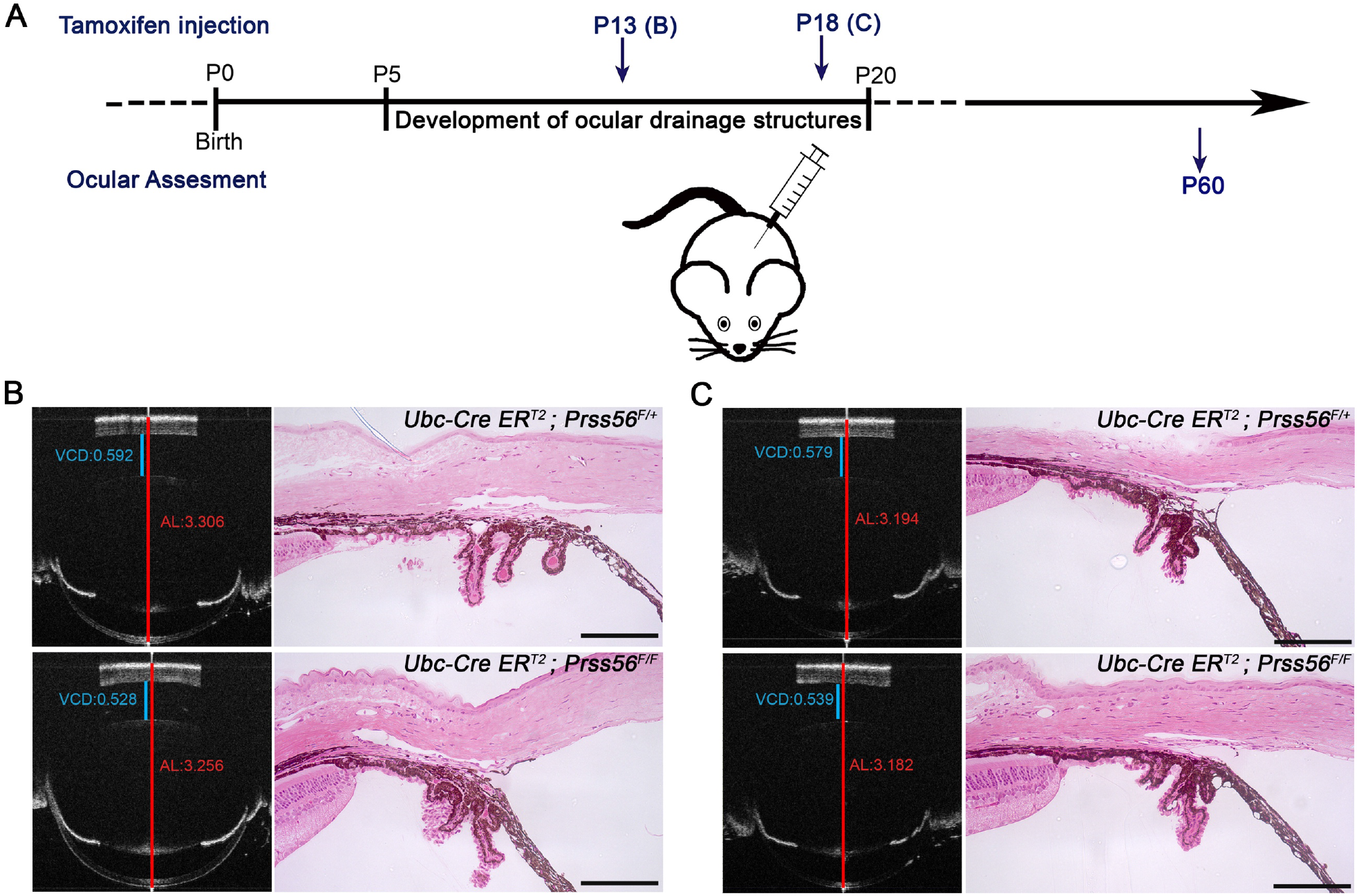
Temporal requirement of PRSS56 for proper positioning of ocular angle structures. To determine the temporal window critical for PRSS56-mediated organization of ocular angle structures, conditional *Prss56* mutant mice (*Prss56^F/F^*) were crossed to mice expressing the ubiquitous inducible Ubc-Cre recombinase (*Ubc-Cre^ERT2^*). Tamoxifen was injected at two different time points to conditionally ablate *Prss56* function at two distinct time points at a stage when the development of the ocular drainage structures is very active (P13) or shortly precedes the completion of their development (P18). (**A**) Schematic diagram illustrating tamoxifen injection paradigm to conditionally ablate *Prss56* function at P13 and P18. (**B-C**) Representative OCT images and H&E stained ocular sections showing significantly reduced ocular axial length and abnormal organization of ocular angle structures leading to angle closure in *Prss56^F/F^; Ubc-Cre^ERT2^* mice following tamoxifen injection at P13 compared to the control *Prss56^F/+^*; Ubc-Cre^*ERT2*^ mice (**B**). In contrast, only a marginal decrease in ocular axial length and no alteration in ocular angle structures organization (as shown by an open angle configuration) were observed in *Prss56^F/F^; Ubc-Cre^ERT2^* mice following tamoxifen injection at P18 compared to control eyes (**C**).

### Mutant Prss56 induced high IOP is dependent on genetic context

The role of genetic context in modulating the severity and penetrance of disease-causing mutations is well established. To determine if IOP elevation observed in mutant *PRSS56* mice are genetic context-dependent, we evaluated IOP in *Prss56* mutant mice maintained on different inbred strains. Shown are data from two genetically divergent inbred backgrounds: C3H/HeJ and DBA/2J (C3H.BLiA-*Pde6b^+^*/J and DBA/2J-*Gpnmb+*/SjJ, respectively) ^19^ that were differentially susceptible too developing high IOP. We found that both the mean IOP and the proportion of *Prss56* mutant mice developing high IOP (>18mmHg, 3 SD above normal IOP) are significantly greater when maintained on a C3H/HeJ (C3H) background compared to a DBA/2J (D2) background at 3 and 6 months of age (Fig. 4A-B). These findings indicate that the C3H background confers susceptibility to IOP elevation caused by a loss of *Prss56* function mutation (*Prss56^glcr4^)*. Notably, the C3H background was also more permissive to the ocular size reduction resulting from a *Prss56* mutation, as shown by a significantly shorter ocular axial length in *Prss56* mutant mice maintained on C3H background compared to those maintained on a *D2* background (Fig. 4C). The more greatly reduced ocular size may or may not exacerbate the IOP phenotype but further experiments are needed to assess this.

**Figure 4.**
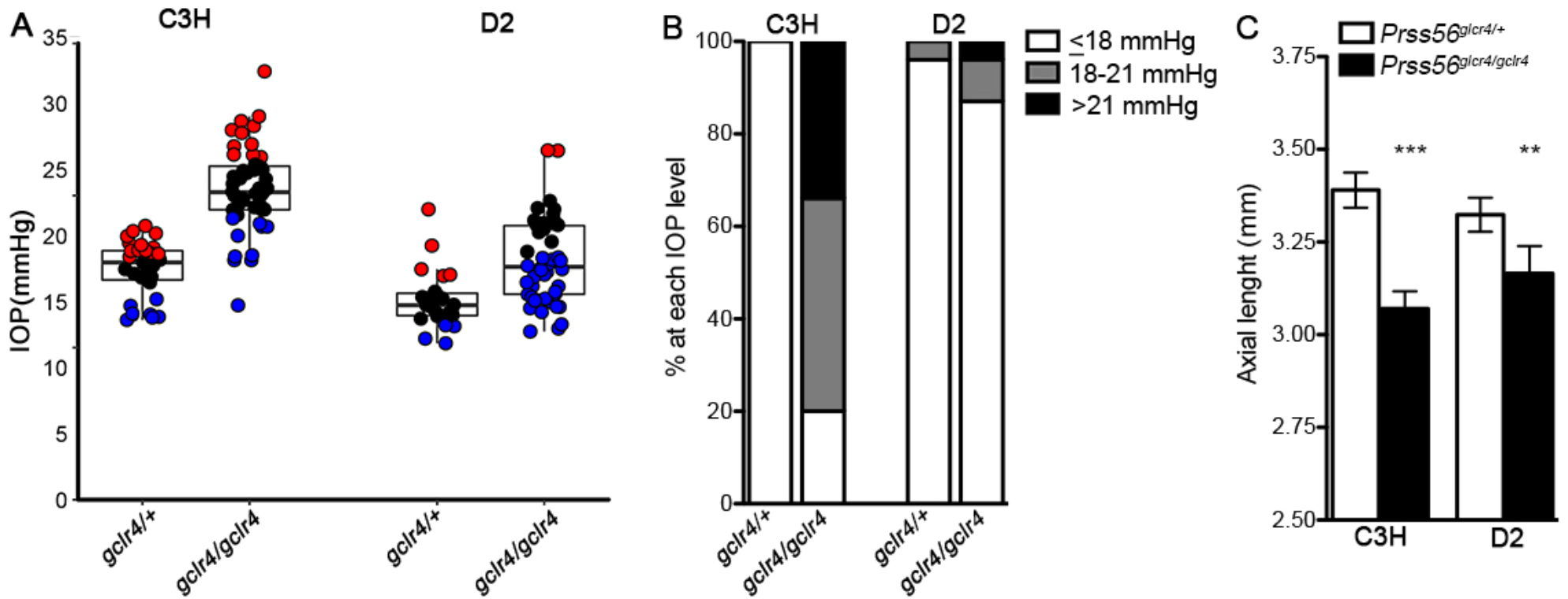
The C3H genetic background is susceptible to high IOP caused by a *Prss56* mutation. (**A**) The *Prss56 ^glcr4^* was backcrossed to C3H and D2 strain over 10 generation to generate respective congenic strains. The IOP values of the congenic *Prss56 ^glcr4^*.C3H and *Prss56 ^glcr4^*.D2 strains and their respective controls is shown as *k-*means clustered (*k*=3) scatter plots. Values are presented as median box plots. The difference in mean IOP of control versus mutant strains was significantly higher on a C3H as compared to the D2 background. (**B**) Percentage of eyes with IOP values at each of binned level at 5 months. (**C**) *Prss56 ^glcr4/glcr4^* mice on a C3H.BliA background also exhibit significant shorter ocular axial length compared to *Prss56^glcr4/glcr4^* mice on a D2 background.

As a first step to identify modifier loci predisposing *Prss56* mutant mice to develop high IOP, we crossed mice that were congenic for the *Prss56* mutation on a C3H genetic background to mice that were congenic for the *Prss56* mutation on a D2 genetic background, which are less susceptible to developing high IOP. The C3HD2F1 progeny was backcrossed to D2.*Prss56* mutant mice and the IOP was measured in the resulting N2 progeny between 3 and 8 months of age. Quantitative Trait Linkage (QTL) analysis was performed to identify genetic regions conferring susceptibility to high IOP caused by the *Prss56* mutation. QTL analysis using IOP values as a continuous trait identified a region on the C3H Chr 2 that is strongly associated with high IOP (Fig. 5A). Furthermore, our analysis shows that this C3H Chr2 locus (60-130 Mb) act in a dominant manner to confer susceptibility to high IOP in response to loss of PRSS56 function (Fig. 5B, Fig.S1).

**Figure 5.**
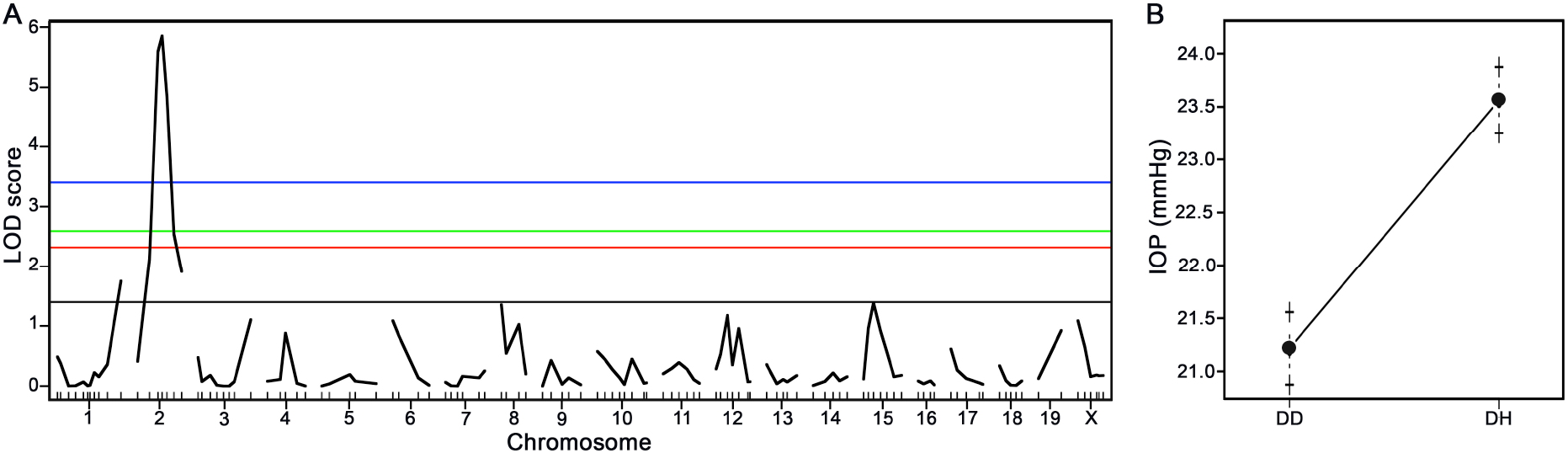
Identification of a modifier locus on Chr2 of the C3H strain that confers susceptibility to high IOP caused by mutant PRSS56. (**A**) Genome-wide QTL analysis of the IOP values measured from the N2 mapping progeny identify a Chr2 locus associated with elevated IOP. The blue, green and red lines indicate the significance thresholds at 1%, 5%, and 10% respectively. (**B**) The effect plot identifies a dominant C3H locus contributes to N2 progeny being susceptible to developing high IOP.

## Discussion

Here, we demonstrate that in addition to its role in ocular size determination, PRSS56 is required for proper positioning of angle structures during ocular development. We show that loss of PRSS56 function during postnatal development leads to an angle-closure phenotype and high IOP. Importantly, our findings suggest that the angle closure phenotype resulting from loss of PRSS56 function is not dependent on ocular axial length reduction. In addition, we demonstrate that the high IOP phenotype resulting from a *Prss56* mutation is genetic context-dependent and we have identified a dominant modifier locus on Chr2 from the C3H genetic background conferring increased susceptibility to IOP elevation.

We have previously reported that a mutation in *PRSS56* caused nanophthamos, a condition characterized by severe reduction in ocular axial length, in humans ^14^. Nanophthalmic individuals are anatomically predisposed to develop a narrow angle and exhibit a high prevalence of angle closure glaucoma^14,20^. Here, we used genetic mouse models to demonstrate that in addition to causing ocular size reduction, loss of PRSS56 function results in altered positioning of the ocular drainage structures and high IOP. Despite nanophthalmic individuals being predisposed to narrow angles, not all of them develop angle closure/high IOP suggesting that reduced ocular size alone is not sufficient to induce angle closure/high IOP^11^. Our data suggest that genetic alterations in factors regulating the positioning of ocular drainage structure during development may predispose individuals to angle closure. In humans, iridocorneal angle configuration is highly variable with a fully open or closed angle at opposite ends of the spectrum. It is tempting to speculate that natural variation in the anatomical configuration of the iridocorneal angle has a genetic basis and influences AqH outflow and susceptibility to developing high IOP.

To address whether ocular axial length reduction contributes to the angle closure and high IOP phenotypes observed in *Prss56* mutant mice, we took advantage of a genetic mouse model (*Egr1^-/-^; Prss56^-/-^*) that rescues the ocular size reduction caused by loss of PRSS56 function^15^. EGR1 is a known regulator of ocular growth^21-23^ and our results show that *Egr1* inactivation significantly rescues ocular axial length reduction in *Prss56* mutant mice while having no significant effect on the angle closure and IOP phenotypes. These findings suggest that the role of *Prss56* in the determination of ocular size is uncoupled from its effect on proper configuration of ocular angle structures. Although, ocular axial length is a commonly used indicator to assess eye size, any change in ocular dimension along the equatorial plane may impact tissue organization. The current state of the art SD-OCT is only capable of precisely measuring ocular length in the axial but not the equatorial plane. A limitation of our study is that we have not assessed the effect of *Egr1* inactivation on the ocular equatorial diameter. Therefore, it remains to be determined if alteration in ocular equatorial diameter can impact the alignment of angle structures.

Ocular expression of *Prss56* is predominantly observed in a subset of Müller glia that is enriched in the peripheral region of the retina^15^. Given the proximity of the peripheral retina to the ocular angle structures, it is possible that PRSS56 secreted from the retina could additionally participate in the development and configuration of ocular angle structures. The formation and positioning of ocular drainage tissues relies on a series of highly orchestrated events starting as early as P0 in the mouse. By P20, drainage structures are fully established as marked by the presence of a well defined Schlemm’s canal and prominent trabecular meshwork layer ^18,25^. Using conditional *Prss56* mutant mice, we demonstrate that the requirement of PRSS56 for normal positioning of the drainage structure coincides with a temporal window critical for the establishment and organization of the ocular drainage tissues ^18^. Being a secreted protease, PRSS56 could contribute to the configuration of ocular angle structures by modulating cell-cell and cell-extracellular matrix interactions.

In both humans with PIC and *Prss56* mutant mice, altered configuration of ocular drainage tissues results in angle closure. However, in contrast to forward displacement of the peripheral iris by anteriorly localized ciliary body in PIC ^12,13,24^, the *Prss56* mutant mice display a posteriorly displaced ocular drainage tissues. In *Prss56* mutant mice, the peripheral iris appears more anteriorly positioned and causes a physical blockage of the ocular drainage tissue. Based on our mouse data, it is tempting to speculate that, in addition to the anterior displacement of the ciliary body and iris, PIC could alternatively result from the posterior displacement of the drainage tissues. This is because both of these conditions result in a similar outcome of aberrant positioning of drainage tissues relative to the iris and ciliary body. Overall, our data suggests that PRSS56 is a critical component of a developmental program that supports proper iridocorneal angle configuration and that alterations in this process may constitute risk factors for ACG.

PACG is generally regarded as a multifactorial complex disorder. Highly heritable ocular anatomical features, including reduced axial length, hyperopia, a relatively larger lens, and a shallow anterior chamber constitute major risk factors for angle closure^26^. In agreement with this underlying complexity, recent GWAS studies have identified several genes associated with PACG^27,28^. Importantly, the identification of modifier genes can provide important and novel mechanistic insights into the ACG pathogenesis. Based on the “Mouse Phenome Database”, there are 292 nonsynonymous coding variants in 765 protein coding genes that are different between the C3H and D2 strain within the confidence interval of the Chr2 locus. The development of key genetic resources and determination of modifier locus lays the foundation for future studies aimed at identification of gene/pathway participating in the pathogenesis of high IOP.

In summary, in this study we have demonstrated that loss of PRSS56 function leads to abnormal positioning of drainage structure leading to their physical obstruction by the peripheral iris, and thereby resulting in angle closure. Thus, we have uncovered a novel role for PRSS56, a serine protease in normal positioning of drainage structure, a feature essential for maintaining an iridocorneal angle in its open configuration. Additionally, we have identified a modifier locus conferring susceptibility to high IOP caused by loss of PRSS56 function. These finding not only provide novel insight into the pathogenesis of ACG but also have significant implication to further our understanding of this devastating blinding disease.

## Method

### Animals

All experiments were conducted in compliance with protocols approved by the Institutional Animal Care and Use Committee at University of California San Francisco (Approval numbers: AN153083 and AN120008) or The Jackson Laboratory’s Institutional Animal Care and Use Committee. Animals were given access to food and water ad libitum and were housed under controlled conditions including a 12-h light/dark cycle in accordance with the National Institutes of Health guidelines.

#### Mouse lines used in this study

*Prss56^glcr4^* mutation: C57BL/6.Cg-*Prss56* ^*glcr4*^/SjJ mice carry an ENU-induced truncation mutation in *Prss56* (protein lacking terminal 104 amino acid) ^14^, *Prss56^Cre^* mutation: C57BL/6.Cg-*Prss56^tm(cre)^* mice have a targeted mutation of *Prss56* in which exon1 was replaced by a sequence coding for CRE recombinase to generate a null allele and induce *Prss56* promoter-driven CRE expression ^33^.

*Prss56* conditional mutation (*Prss56^F^*): C57BL/6.Cg-*Prss56^tm1^/*SjJ mice carry LoxP sites flanking exons 2 to 4 of *Prss56*, which result in a catalytically inactive protease in a Cre-dependent manner.

*Egr1* mutation: B6;129-*Egr1^tm1Jmi^*/J mice have a targeted mutation of *Egr1* resulting in a null allele.

*Ubc-Cre ER^T2^* transgene: C57BL/6.Cg-Tg(UBC-Cre/ER^T2^)1Ejb mice express the ubiquitous inducible Ubc-Cre recombinase (*Ubc-Cre^ERT2^*) in response to tamoxifen.

### IOP Measurement

IOP was measured using the microneedle method as described previously ^34^. In brief, the mice were anesthetized by intraperitoneal injection of a mixture of ketamine (99 mg/kg; Ketalar, Parke-Davis, Paramus, NJ) and xylazine (9 mg/kg; Rompun, Phoenix Pharmaceutical, St. Joseph, MO). IOP was measured in both male and female mice. Prior to measuring IOP on experimental mice, the device was calibrated. We measured IOP in a small cohort of C57BL/6J mice along with the experimental mice as a methodological control. To show the difference in the number of mice with elevated IOP in each group we displayed the data using *k*-means clustering (*k*=3).

### Ocular biometry

We performed mouse ocular biometry using the Envisu R4300 spectral domain optical coherence tomography (SD-OCT, Leica/Bioptigen Inc., Research Triangle Park, NC, USA) as described previously ^35^. Mice were anesthetized with ketamine/xylazine (100 mg/kg and 5mg/kg) and their eyes dilated before performing ocular biometry measurements. The distance from the corneal surface to the retinal pigment epithelium/choroid interface provided axial length measurements. VCD was calculated by measuring the distance between the innermost layer of the retina and the lens. To minimize any confounding effect of body weight on ocular size, we only selected littermates with comparable body weights in each of the experimental groups.

### Ocular Histology

After euthanizing the mice, their eyes were enucleated and immersed in cold fixative (1% PFA, 2% glutaraldehyde, and 0.1 M cacodylate buffer cacodylate buffer) for a minimum of 48 hours. The fixed eyes were then transferred to 0.1 M cacodylate buffer solution and stored in a refrigerator till further use. The eyes were then embedded in glycol methacrylate, and then sectioned using a microtome. Serial sagittal sections (2μm) were made by passing through the optic nerve and the sections were stained with hematoxylin and eosin (H&E). Hematoxylin and eosin stained ocular sections were assessed to determine the configuration of ocular angle structures. We evaluated at least 15-30 similarly spaced sections from 2 ocular regions (mid-peripheral and central).

### Gene Mapping and QTL studies

The *Prss56* mutant allele (*Prss56 ^glcr4^*) was introduce into the two genetically divergent C3H/HeJ (C3H) and DBA/2J (D2) inbred mouse strains and backcrossed for at least 10 generations. D2 mice spontaneously develop high IOP and glaucoma due to a mutation in *Gpnmb*. Hence, we have used the DBA/2J-*Gpnmb^+^*/SjJ sub-strain of D2 mice that has been corrected for the mutation in *Gpnmb*. Similarly, we utilized a C3H substrain corrected for the inherent *PDE6b* mutation causing retinal degeneration (C3A.BLiA-*Pde6b*^+^/J). To identify the genetic locus predisposing *Prss56* mutant mice to develop high IOP on the C3H background, *Prss56^glcr^4*.C3H mice were bred to D2 mice, which are typically less permissive to developing high IOP and the resulting *Prss56^glcr4^*CH3.D2 F1 progeny was backcrossed to the D2 background. A total of 175 N2 progeny mice were aged up to 9 months and their IOP was measured at multiple time points between the ages of 3 to 9 months. Genotyping was performed using over 116 genome-wide single nucleotide polymorphic markers that are distinct between the C3H and D2 genomes and spaced at approximately 20 MB intervals (SNPs, KBioscience, UK). Genotyping quality assessment was done by evaluating the recombination fraction plot and map plot ^36^. Pseudo-markers were generated at 2 cM spacing for each chromosome. The log transformed IOP values showed a normal distribution were therefore used for further analysis (Fig S1). We performed a genome-wide one- QTL scan using 256 imputations to identify the chromosomal loci regulating IOP (NCBI Build 37). Thousand permutations were performed to calculate the three thresholds for QTL detection at 1%, 5%, and 10%. A QTL with LOD score above the 1% threshold was identified as a strong QTL, while anything at or above 10% were suggestive QTL.

## Acknowledgements

Yien-Ming Kuo for histology and staining. This work was supported in part by NIH P30 core grant for vision research (UCSF, Ophthalmology), Research to Prevent Blindness unrestricted grant (UCSF, Ophthalmology) and William and Mary Greve Special Scholar Award (KSN), That Man May See Inc (KSN), Research Evaluation and Allocation Committee (REAC)-Tidemann fund (KSN), Marin Community Foundation-Kathlyn McPherson Masneri and Arno P. Masneri Fund (KSN). Knight Templar Eye Foundation Career Starter Award (SP) and NEI grants EY022891 (KSN), EY011721 (SWMJ).

## Author’s contributions

CL-D, NT, SP, SWMJ and KSN conceived and performed experiments, analyzed the data and prepared the manuscript. KSN oversaw all aspects of the study. YS and ED contributed towards colony management, genotyping and histological analysis.

---

**Fig S1.**
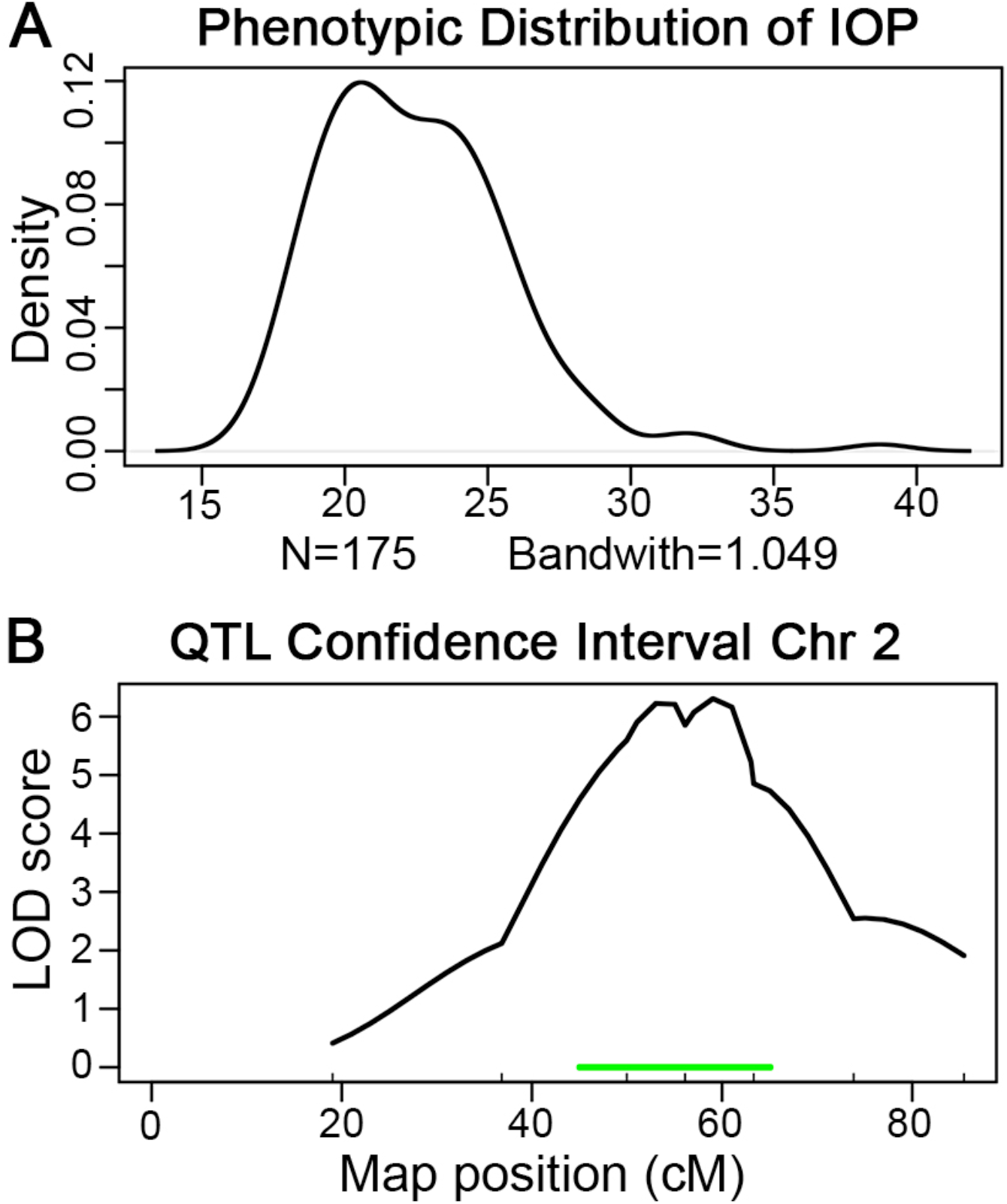
**A.** Plot showing log transformed IOP values showing a normal distribution. **B.** Plot showing QTL confidence interval on chromosome 2.

